# Periodized plyometric training improves jumping performance and reduces force-velocity imbalance in collegiate volleyball players

**DOI:** 10.64898/2026.07.22.740194

**Authors:** Hong-Mei Wu, Xue-Feng Xi, Jun Li

**Affiliations:** School of Physical Education, Xinyang Agriculture and Forestry University, Xinyang, Henan 464000, China; School of Physical Education, Henan University, Kaifeng, Henan 475001, China

**Keywords:** volleyball, plyometric exercise, force-velocity relationship, countermovement jump, training periodization

## Abstract

Few studies have focused on plyometric interventions lasting over 24 weeks in team-sport athletes, and little research has validated the practical utility of personalized force-velocity assessment. A total of 36 university volleyball athletes (21.35±1.87 years) were randomly allocated into either a 32-week plyometric training group (PT, n=18) or a low-intensity active control group (CON, n=18); 34 players completed the trial (17 per group). PT completed three weekly training sessions, while CON maintained routine technical practice and general low-intensity conditioning. CMJ height, RSI_mod_, and force-velocity profiles (F₀, V₀, peak power, FV imbalance) were evaluated at baseline and at weeks 8, 16, 24, and 32. PT increased CMJ height from 33.1±4.8 cm to 38.4±5.2 cm, a net gain of 5.3 cm (p<0.01, d=0.99, 95% CI [0.57, 1.41]); CON showed negligible change (+0.5 cm, p=0.68). The PT group also improved RSI_mod_ by 31.6% (d=0.87, 95% CI [0.45, 1.29], p<0.01) and peak power by 18.4% (d=0.76, 95% CI [0.35, 1.17], p=0.004). Meanwhile, FV imbalance dropped markedly by 31.4%, from 28.3% to 19.4% (d=–0.68, 95% CI [–1.09, –0.27], p=0.024). No significant F-V changes occurred in CON. The reduction in FV imbalance correlated moderately with CMJ gain (r= – 0.53, p=0.024), whereas the change in peak power did not (r=0.21, p=0.39). Overall, 32 weeks of periodized plyometric training enhanced jump performance, reactive strength, and power, and reduced the force-velocity (FV) imbalance by approximately 31%. The magnitude of improvements in jump performance (16%) and FV imbalance reduction (31%) observed in this 32-week periodized program fall within a similar range to previously reported effects of individualized F-V training (14% and 40%, respectively). However, direct numerical comparisons are confounded by differences in study design, population, and methodology; head-to-head trials are needed.

## INTRODUCTION

Volleyball requires athletes to jump explosively many times per match. In five-set college matches, middle blockers perform about 118 jumps on average [1]. Consequently, developing lower-limb power is a central goal for coaches. Plyometric training has been a cornerstone of power development for decades. A 2019 systematic review by Silva and co-workers [2] concluded that PT improves vertical jump in volleyball players, but most of the included studies lasted only 6–12 weeks – much shorter than a full competitive season. Whether extending PT to 32 weeks yields additive benefits or merely maintains earlier gains remains unknown.

The force-velocity (F-V) relationship provides a mechanical framework for understanding and evaluating jumping ability. From squat jumps performed with increasing loads, the F-V relationship can be derived, yielding estimates of theoretical maximal force (F₀), maximal velocity (v₀), maximal power (Pmax), and an imbalance index (FVimb) that identifies force-deficient or velocity-deficient athletes [3]. Individualising training based on F-V profiling has been shown to improve jump height more effectively than generic programs [4,5]. However, such individualised prescription requires laboratory testing and per-athlete programming, which is often impractical for university teams.

Currently, the efficacy of a team-based plyometric program in modifying the force–velocity (F-V) profile and reducing FV_imb_ remains undetermined. Previous investigations have typically adopted one of two approaches: either they were confined to short intervention periods (≤ 9 weeks), or they employed highly individualized training prescriptions tailored to each athlete’s F-V characteristics. To address this gap, the present trial was designed to examine whether a standardized, periodized plyometric protocol delivered to an entire team could produce meaningful alterations in lower-limb F-V parameters. No trial has used a 32-week periodized PT protocol in volleyball players with serial F-V assessments.

Recent evidence has increasingly focused on F-V-based individualized training. The systematic review and meta-analysis by Solberg et al. [5] synthesized available evidence, and the more recent review by Wolte et al. [6] further indicated that individualized training tailored to an athlete’s F-V profile can partially correct force deficits and fully correct velocity deficits, but confers almost no additional benefit for athletes who already possess an optimal profile. Notably, jump height gains from individualized training were not significantly superior to those achieved with well-designed generic programs. This finding carries important implications for team sports, where individual F-V testing for each athlete is often impractical to implement in real-world settings. Subsequently, Iranpour et al. [7] showed that just four weeks of plyometrics incorporating speed and weight overloads improved both jump performance and explosive power in male volleyball players. Their work suggests that strategic manipulation of training variables—not necessarily individualized diagnosis—can trigger meaningful adaptations. Together, these recent findings support the concept that a carefully periodized, team-based plyometric program can meaningfully improve the F-V profile without the logistical burden of one-on-one testing.

Three primary hypotheses were formulated prior to the intervention. We hypothesized that the 32-week general training plan would elicit measurable gains in CMJ performance, modified RSI and peak power. We also anticipated that force–velocity imbalance would be lowered through this standard approach, without requiring individually tailored training modifications. In addition, changes in CMJ jump ability were expected to follow reductions in FV imbalance more closely than standalone increases in peak power. With these findings in mind, the current trial asked whether a 32-week periodized plyometric program—delivered uniformly to an entire volleyball team—could alter lower-limb force–velocity characteristics and boost jumping ability in collegiate players.

## METHODS

### Study Design and Ethics

This was an assessor-blinded, parallel-group randomised controlled trial, conducted in accordance with the Declaration of Helsinki. The institutional review board granted ethical approval by XYAFU Ethics Committee(No. XY-2024-029), and all participants provided written informed consent prior to enrolment.

### Participants

Participants were recruited and screened for eligibility between 01/01/2024 and 31/03/2024 (a three-month period). All participants were recruited from the first-team rosters of the university’s men’s and women’s volleyball programs. A total of 36 athletes—18 men and 18 women—agreed to take part and provided written informed consent before any testing or training commenced. Their mean age was 21.35 years (standard deviation 1.87 years), and all were actively competing at the collegiate level.

Four inclusion criteria were applied. Being an official team member for at least 12 months before the study started was mandatory, which ensured a basic level of volleyball-specific conditioning and tactical understanding. In addition, no lower-limb injury—including ankle sprains, knee ligament damage, or muscle strains—could have occurred in the six months before recruitment. This rule lowered the chance of re-injury during high-impact plyometric work and avoided confusion that might arise from incomplete healing. Participants also could not be doing any systematic plyometric training outside their regular team practices; this guaranteed that any training effect measured would come solely from the prescribed 32-week program, not from unsupervised extra jumping. Finally, each athlete had to keep a weekly volleyball practice volume of at least eight hours—a typical load for collegiate players during the competitive season—so that everyone started from a comparable baseline of sport-specific loading.

Exclusion criteria were also applied. We excluded anyone who had been diagnosed with cardiovascular disease (e.g., arrhythmias, uncontrolled hypertension) or metabolic disorders (e.g., type 1 or type 2 diabetes) that could be exacerbated by intense plyometric exercise. We also excluded individuals who had undergone any surgical procedure on the lower limbs—including but not limited to anterior cruciate ligament reconstruction, meniscal repair, or ankle stabilization surgery—within the three months preceding the study. Finally, we did not allow enrollment of athletes who were currently using anabolic steroids or systemic corticosteroids, as these substances can influence muscle force production, recovery capacity, and tendon integrity, thereby confounding the interpretation of training-induced adaptations. No participants were excluded based on these medical or pharmacological criteria, and all 36 recruited athletes successfully passed the screening process.

### Sample Size Calculation

We estimated the required sample size using G*Power (version 3.1) for a two-tailed independent-samples t-test (α = 0.05, power = 0.80). Based on the sub-group analysis of athletes aged ≥16 years reported in a previous meta-analysis of plyometric training in volleyball players (ES = 1.28; Ramirez-Campillo et al., 2021 [8]), the calculation yielded a minimum of 11 athletes per group. To be more conservative and to account for an anticipated attrition rate of approximately 15% over the 32-week intervention, we inflated this number, targeting 18 participants per group (36 total at baseline). By the end of the program, we had lost only two players (5.6%), leaving 34 completers (17 per group). A CONSORT flow diagram is provided as Supplementary Figure S1.

### Randomisation and Blinding

To reduce selection bias and ensure baseline comparability, the following procedures were implemented. A statistician independent of recruitment and testing generated the allocation sequence using a computer random number generator. To ensure each group had a similar mix of men and women — given possible sex differences in baseline jump ability — we stratified randomization by sex. Within each sex stratum, we used block randomization with a fixed block size of four. This approach kept the two groups numerically balanced during recruitment without making the next assignment guessable.

For allocation concealment, we used sequentially numbered, opaque, sealed envelopes. Each envelope held the group assignment for a single participant, following the pre-generated sequence. The same statistician prepared the envelopes and kept them in a locked cabinet. Envelopes were opened only after a participant had completed all baseline tests, including CMJ and force-velocity profiling. As a result, neither the research team nor the participants knew the group assignment before baseline measurements finished.

We implemented an assessor-blinded design to minimize detection bias. The lead testing technician, who conducted all outcome assessments (CMJ, RSImod, F-V parameters) at baseline and at weeks 8, 16, 24, and 32, was blinded to group allocation and had no access to training schedules. The analyst performing the final statistical analysis was also blinded until the database was locked and all quality checks were completed. Group codes were revealed only after the pre-specified statistical analysis plan had been executed.

Participant and coach blinding was not feasible due to the inherently different nature of the interventions (high-intensity plyometric training versus low-intensity conditioning). However, participants were instructed not to discuss their training content with the testing technician, and all testing sessions were conducted in a standardized environment with minimal interaction between participants and the assessor.

### Training Interventions

#### Plyometric group (PT)

The PT group followed a 32-week periodized program that replaced some general conditioning within their regular volleyball practice. Sessions ran three times weekly, each lasting 45–60 minutes, with progression based on technical mastery. The program had four phases:

Weeks 1–8 (anatomical adaptation): low-impact drills including ankle hops, pogo jumps, and box drops from 15–25 cm. Contact volume: 80–120 per session.

Weeks 9–16 (strength-power transition): hurdle hops, drop jumps from 30–45 cm, and vest-loaded jumps (5% body mass). Contacts: 100–150 per session.

Weeks 17–24 (maximal power): depth jumps (50–70 cm), 60 cm box jumps, and squat jumps with a 10% body-mass vest. Volume lowered to 60–80 contacts to keep intensity high.

Weeks 25–32 (maintenance): two sessions weekly, mixing low- and high-intensity drills, 40–60 contacts per session.

A certified strength and conditioning coach watched over all sessions. Athletes moved to a higher load only when they could do the current level with good form.

#### Control group (CON)

The CON group did regular team drills plus low-intensity conditioning work – core stability, resistance bands, balance exercises – three times a week, 45–60 minutes per session. This kept weekly supplementary training time the same as in the PT group, though the mechanical stress was very different.

### Testing Procedures

Assessments took place at baseline (week 0) and after each phase (weeks 8, 16, 24, 32). Each test session was scheduled 48 h after the last training bout to avoid acute fatigue. All sessions were conducted in the same biomechanics laboratory (temperature 22 ± 1°C, humidity 50%).

Warm-up: 5 min cycling at 50 W, followed by 3 min dynamic stretching and 3 submaximal CMJs.

Force plate setup: Two AMTI BP600900 plates (1200 Hz) embedded flush with the floor. Signals were low-pass filtered (Butterworth, 4th order, cutoff 50 Hz). Vertical ground reaction force (vGRF) was used to compute flight time, jump height, and propulsion velocity.

CMJ protocol: Participants stood upright with hands on hips, performed a rapid countermovement to a depth corresponding to a knee angle of approximately 90° (range: 80–100°), then jumped maximally without arm swing. Knee angle at the lowest position was verified offline from sagittal-plane video recordings (sampled at 60 Hz); trials falling outside the prescribed range were discarded and repeated. Three valid trials (≤5% variability in height) were recorded; the trial with the highest jump height was used for analysis. Trials with visible countermovement hesitation or incomplete extension at take-off were discarded and repeated. Each trial was separated by 30 s of passive rest, with a 3-min rest interval between sets of jumps.

Reactive Strength Index modified (RSI_mod_): Following Ebben and Petushek[9], RSI_mod_ was calculated from the CMJ as jump height (m) divided by time to take-off (s), where time to take-off was defined as the interval from the onset of the countermovement (first detectable vertical movement) to the instant of take-off.

Force-velocity (F-V) profile:The two-point method described by Samozino et al.[10] was used. Participants performed maximal squat jumps (without countermovement, hands on hips) under two conditions: unloaded (body mass only) and with a weighted vest adding 20% of body mass (BM). Three attempts per condition were recorded; the jump with the highest flight-time-derived height was retained. From the linear regression F = F₀ – S·v (where S is the slope), theoretical maximal force (F₀) and velocity (v₀) were estimated. Maximal power (P_max_) was calculated as F₀·v₀/4 (expressed in W/kg). The force-velocity imbalance (FV_imb_) was computed following Morin and Samozino [11]:FV_imb_(%) = (v₀_actual_ – v₀_opt_) / v₀_opt_ × 100 where v₀_opt_ is the optimal velocit y that would yield the same P_max_ under a balanced F-V slope. The two-point method has been shown to provide reliable estimates of F₀, v₀, and P_max_[12]. In a preliminary test-retest session (n=17, 48 h apart), we observed ICC >0.90 for F₀ and P_max_, and TEM <5% for all F-V variables; the minimal detectable change (MDC_95_) for FV_imb_ was 3.8%. The choice of 20% BM was based on pilot testing (n=17), which identified it as the transition point between force- and velocity-dominant regions; loads in the 20–30% BM range provide reliable F-V slope estimates with low injury r isk [13] and reduce testing duration and fatigue.

### Statistical Analysis

All values are shown as mean ± SD, where SD denotes the standard deviation.The primary outcome was CMJ height measured at week 32; secondary outcomes included RSI_mod_, P_max_, and FV_imb_. Statistical analyses employed a mixed-model repeated-measures ANOVA, incorporating fixed effects for group (PT vs. CON), time (five assessment points), and their interaction, along with a random intercept for each participant. Model fitting was carried out using the lmer function from the lme4 package in R. Kenward-Roger degrees of freedom were applied. Post-hoc pairwise comparisons were adjusted using Bonferroni correction (10 comparisons; within-PT threshold α=0.005). Effect sizes were Cohen’s d with 95% confidence intervals derived from the non-central t-distribution. Pearson correlations were calculated between change scores (Δ = week 32 – baseline) for CMJ and F-V parameters. Missing data (2.8% of total data points, due to illness on test days) were handled by restricted maximum likelihood (REML). Statistical significance was set at two-tailed α = 0.05. All analyses were performed in R 4.2.2 and SPSS 26.0. For visualization purposes in Figure 5A, participants were categorized by baseline FVimb using the median split of the PT group distribution (≥28% vs. <28%); these subgroup comparisons were exploratory and not pre-specified.

To account for potential sex differences arising from the stratified randomisation, sex was included as a fixed covariate in an additional mixed-model analysis; the results are presented in Supplementary Table S2.

### Monitoring of training load and confounding factors

Throughout the 32-week intervention, we monitored several variables to characterise the external and internal training load of both groups. Volleyball technical/tactical training time (hours per week) and match exposure (number of sets played per week) were recorded via team logs. Resistance training outside the prescribed protocol was prohibited; compliance was checked by weekly self-report questionnaires. Perceived exertion (session-RPE, 0–10 scale) was collected for each PT and CON session to quantify internal load. Injury and illness episodes were documented by the team physiotherapist. For female participants (n=18), menstrual cycle phase at each test session (baseline, weeks 8, 16, 24, 32) was recorded via self-report, and any use of oral contraceptives was noted. Sleep duration (hours per night) and subjective sleep quality (1–5 scale) were assessed weekly using a short diary. Nutritional intake was not standardised, but participants were instructed to maintain their habitual diet; no ergogenic supplements were permitted.

No significant differences between PT and CON groups were observed for any of these monitored variables (all p > 0.05; see Supplementary Table S1 for full data). To ensure that the additional jumping load from regular volleyball technical drills did not differ between groups, we estimated the number of jumps performed during routine practice sessions. These data are summarised in Supplementary Table S3. Therefore, the observed between-group differences in primary outcomes can be reasonably attributed to the plyometric intervention rather than to differential exposure to other training or lifestyle factors. Nevertheless, the absence of continuous objective monitoring (e.g., GPS-based external load, heart rate-based internal load) limits the precision of this adjustment.

## Results

### Participant Flow and Baseline Characteristics

Two participants withdrew: one in PT (ankle sprain during a match, unrelated to the intervention) and one in CON (academic scheduling conflict). Thus, 34 players completed the trial (PT n=17, CON n=17). Baseline characteristics were well balanced between groups (Table 1). Adherence to PT sessions averaged 95.2% (range 88–100%).

**Table 1.** Baseline characteristics of completers.

| Variable | PT (n=17) | CON (n=17) | p |
| --- | --- | --- | --- |
| Age (yr) | 21.4 ± 1.9 | 21.2 ± 1.7 | 0.74 |
| Height (cm) | 183.5 ± 8.2 | 184.1 ± 7.9 | 0.83 |
| Body mass (kg) | 76.2 ± 9.5 | 75.8 ± 10.1 | 0.91 |
| Training experience (yr) | 6.6 ± 1.8 | 6.4 ± 2.0 | 0.76 |
| Sex (M/F) | 9/8 | 8/9 | 0.73 |

### CMJ Height and Reactive Strength

A significant group × time interaction emerged for CMJ height (F_4,128_ = 9.73, p < 0.001, ηp² = 0.233). As shown in Figure 2A, PT progressively increased CMJ from week 16 onward, reaching a gain of 5.3 cm at week 32 (95% CI 3.0 to 7.6; d = 0.99, p < 0.01). CON showed only a trivial change (Δ = +0.5 cm, p = 0.68). Between-group differences became statistically significant at week 16 and remained so thereafter.

**Figure 1.**
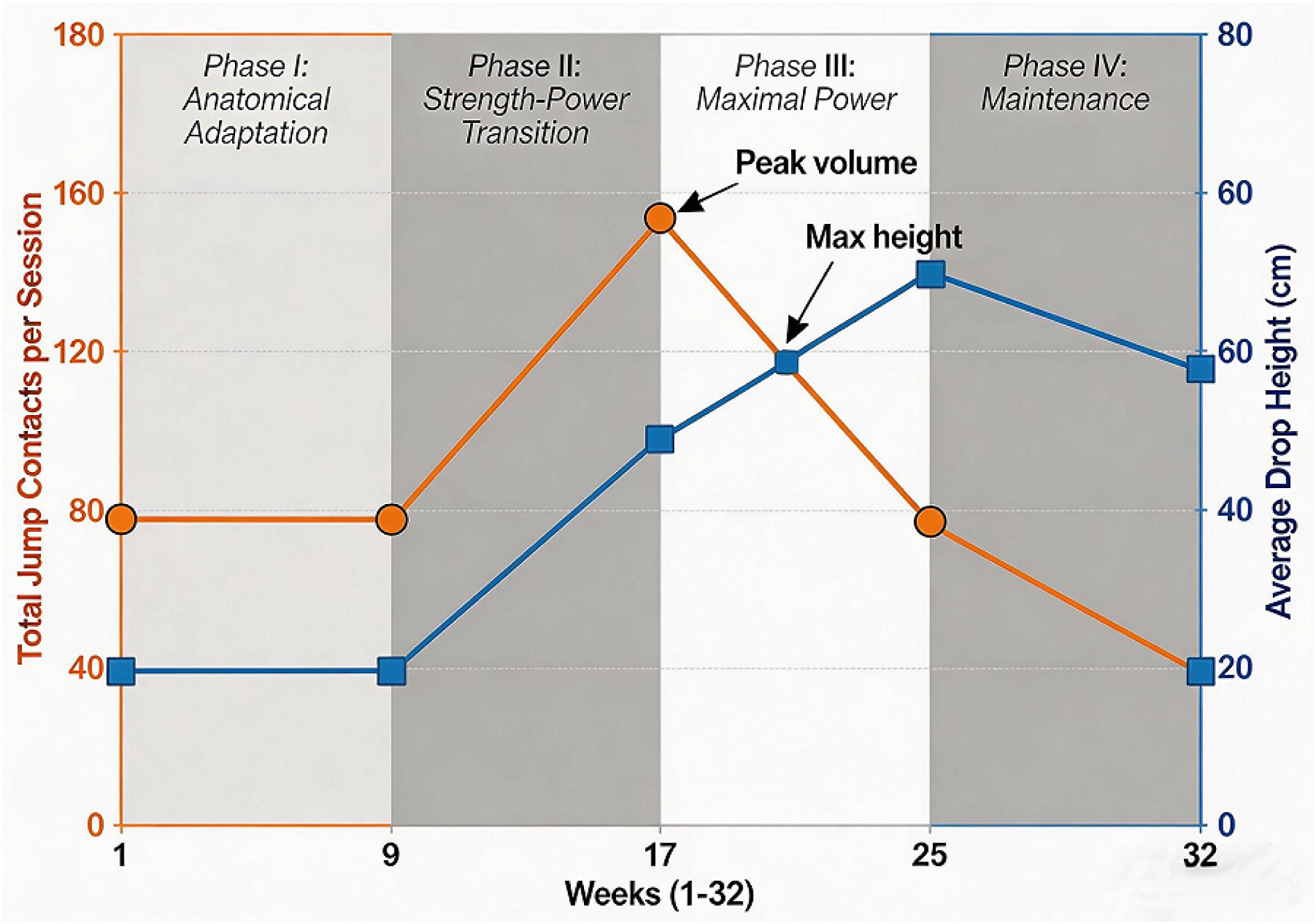
Weekly training load progression across the 32-week periodized plyometric program. Orange line (left Y-axis, circles): jump contacts per session; blue line (right y-axis, squares): average box/drop height (cm). Data points are linearly interpolated between control weeks 1, 9, 17, 25, and 33. Shaded vertical bands (alternating light and medium grey) indicate the four training phases: I – anatomical adaptation (weeks 1-8), II – strength-power transition (weeks 9-16), III – maximal power (weeks 17-24), IV – maintenance (weeks 25-32). Phase labels are centered at the top (y=170). Arrows highlight the peak contact volume (150 at week 16-17) and the maximum drop height (65 cm at week 24). The load follows a periodized undulating pattern, peaking in Phase III before reducing during maintenance.

**Figure 2.**
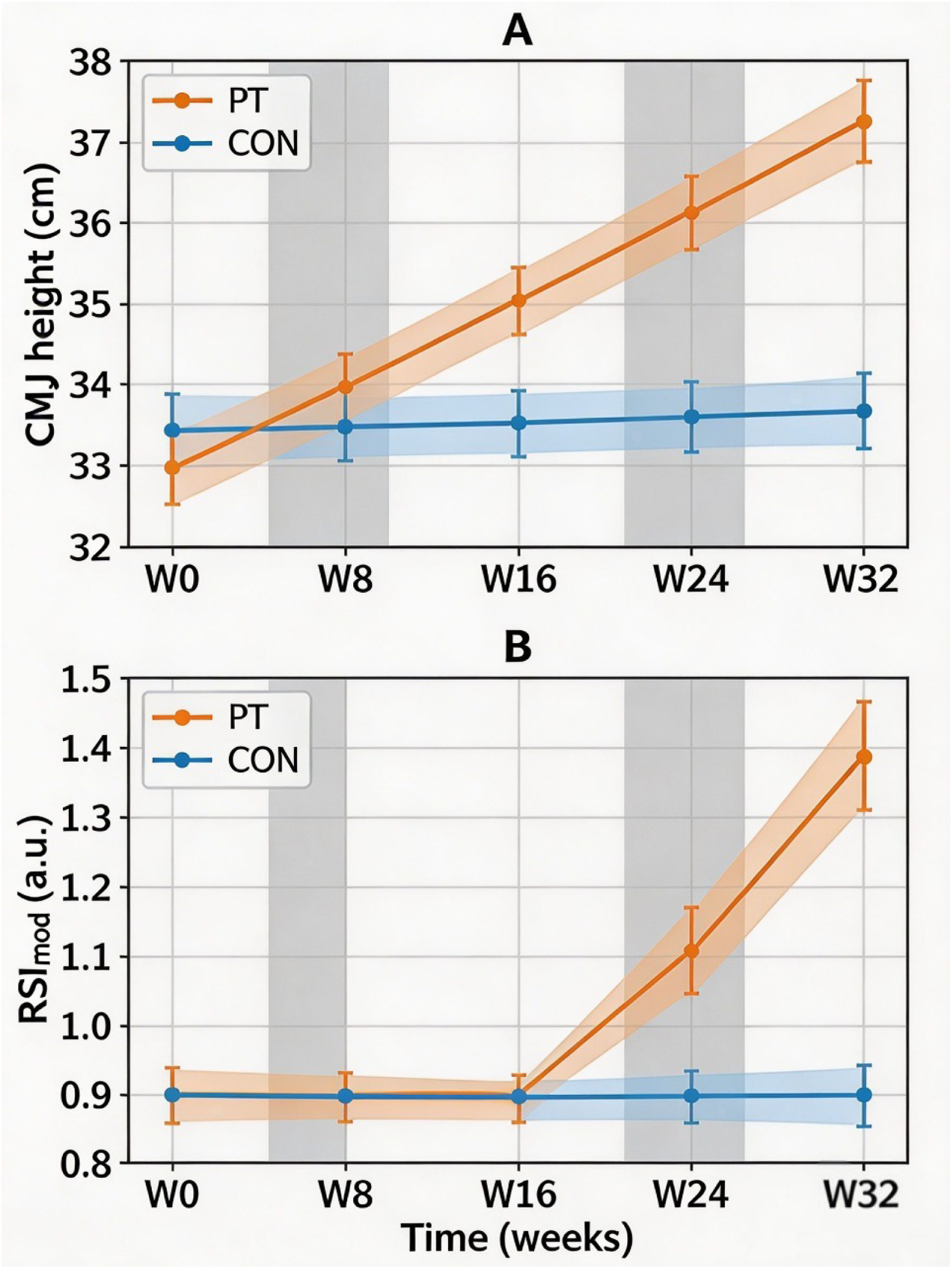
Temporal changes in (A) countermovement jump (CMJ) height and (B) reactive strength index-modified (RSI_mod_) over 32 weeks. Data are presented as mean ± SD. PT: plyometric training group (n = 17); CON: control group (n = 17). Vertical shaded bands denote the four training phases: Phase I (weeks 1–8), Phase II (weeks 9–16), Phase III (weeks 17–24), Phase IV (weeks 25–32). Significant group × time interactions were observed for both outcomes (mixed-model ANOVA: both p < 0.001). Significant within-PT improvements (p < 0.005, Bonferroni) are indicated by asterisks (*) from week 16 onward. No significant changes occurred in CON.

RSI_mod_ also exhibited a significant group × time interaction (F_4,128_ = 5.21, ηp² = 0.140). PT improved from 0.41 ± 0.09 to 0.54 ± 0.11 (+31.6%, d = 0.87) by week 32, whereas CON remained unchanged (0.42 → 0.43, p = 0.75).

**Figure 2** presents the evolution of CMJ height (panel A) and RSI_mod_ (panel B) across the 32 weeks. Data are means ± SD. Shaded vertical bands mark the four training phases. The PT group shows a clear upward trend from week 16 onward, while CON lines remain flat.

**Figure 3A** (violin plot with boxplot) compares the distribution of week 32 CMJ values between groups. PT participants had a higher median, whereas the spread (IQR) was comparable between groups. **Figure 3B** (spaghetti plot) displays individual CMJ trajectories: semi-transparent lines represent individual participants, bold lines indicate group means. Nearly all PT participants exhibited a progressive upward shift, whereas CON trajectories stayed essentially flat.

**Figure 3.**
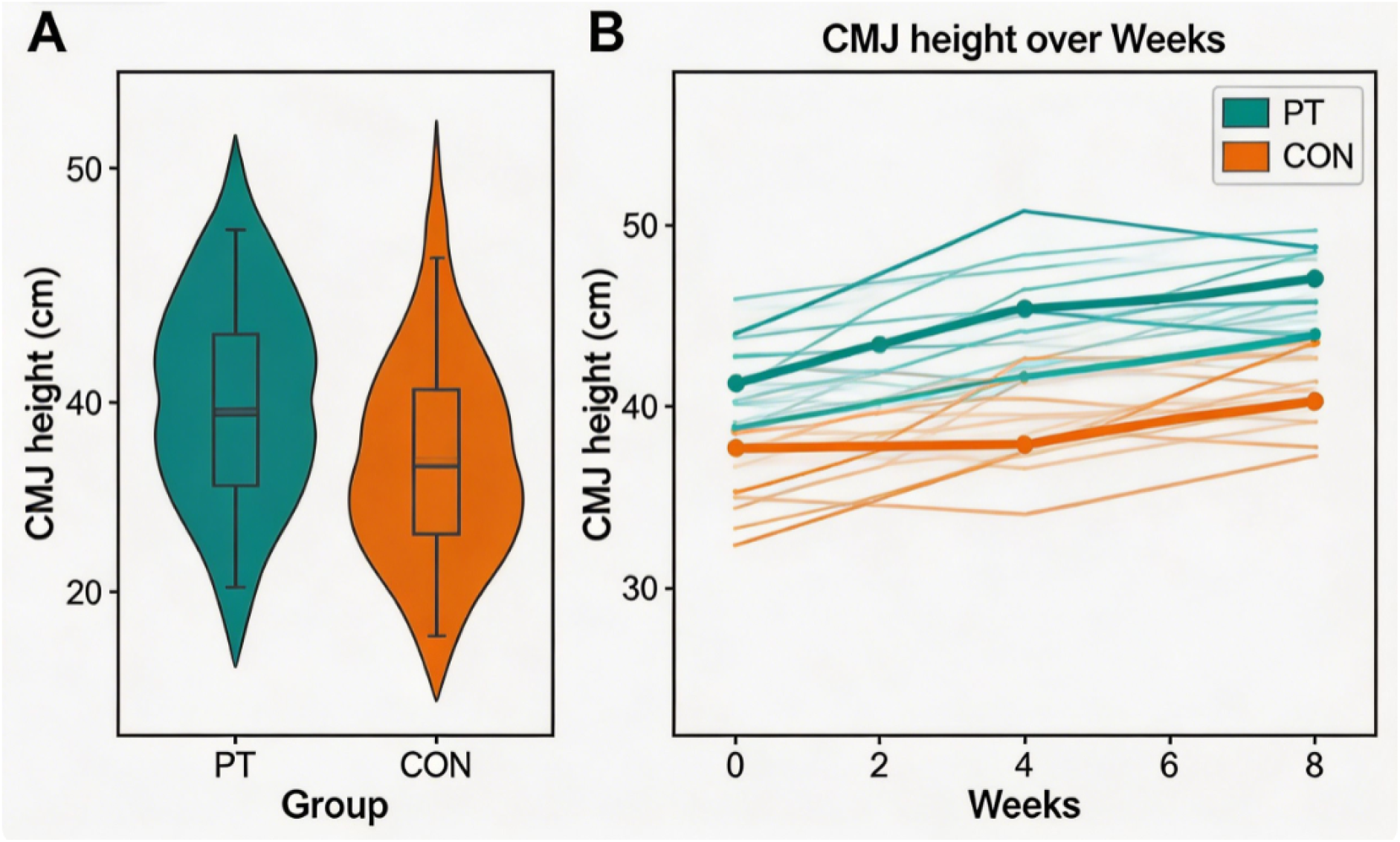
(A) Violin plots with overlaid boxplots showing CMJ height distribution at week 32 for the PT (n=17) and CON (n=17) groups. The white dot represents median; box spans IQR; whiskers extend to 1.5×IQR. Violin width indicates probability density. Corresponding summary statistics for week 32 are presented in Table 2. (B) Individual spaghetti plots of CMJ height over 32 weeks for PT (left) and CON (right). Semi-transparent thin lines represent individual participants; bold solid lines represent group mean trajectories. The group mean values at each time point are reported in Table 2. PT shows a clear progressive upward shift for most individuals, whereas CON trajectories remain nearly flat.

**Table 2.** CMJ height (cm) over time (mean ± SD).

| Time | PT (n=17) | CON (n=17) | Within-PT $\Delta$<br>(95%CI) | p (PT vs. CON) |
| --- | --- | --- | --- | --- |
| Baseline | 33.1 ± 4.8 | 33.6 ± 4.3 | – | 0.67 |
| Week 8 | 34.6 ± 5.0 | 33.8 ± 4.5 | +1.5 (-0.2, 3.2) | 0.18 |
| Week 16 | 36.3 ± 5.1 | 33.9 ± 4.4 | +3.2 (1.0, 5.4)‡ | 0.003 |
| Week 24 | 37.5 ± 5.2 | 34.0 ± 4.5 | +4.4 (2.2, 6.6)‡ | 0.002 |
| Week 32 | 38.4 ± 5.2 | 34.1 ± 4.5 | +5.3 (3.0, 7.6)‡ | 0.004 |

‡ Bonferroni-adjusted $p < 0.005$ vs. baseline within PT.

**Figure 4** presents the individual changes in CMJ height from baseline to week 32. All but one participant (94%) improved beyond the typical error of measurement (TEM = 1.2 cm), indicating a robust response to the intervention. Subgroup analyses by sex are presented in Supplementary Figure S2; both male and female athletes in the PT group exhibited progressive improvements, with no significant sex-by-training interaction (Group × Time × Sex: p = 0.349).

**Figure 4.**
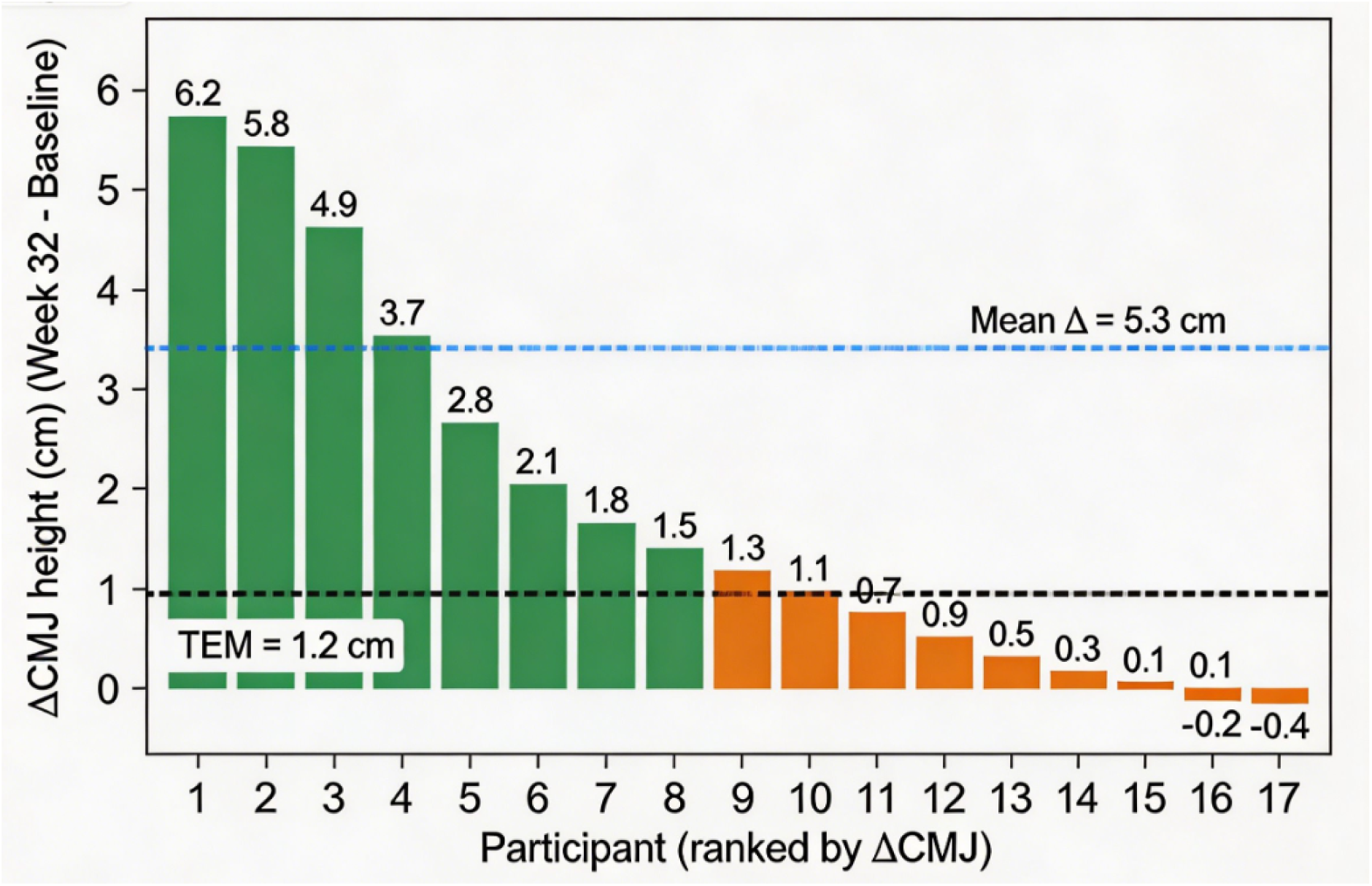
Waterfall plot of individual changes in CMJ height (ΔCMJ) from baseline to week 32 in the plyometric training group (n=17). Participants are sorted from largest to smallest improvement. Error bars represent the typical error of measurement (TEM = 1.2 cm), estimated from test-retest reliability. The dashed horizontal line indicates the TEM. All but one participant (94%) showed improvements exceeding the TEM, indicating a robust response to the intervention.

### Force-Velocity Profile

**Table 3** summarises the F-V parameters. PT increased P_max_ from 24.3 W/kg to 27.5 W/kg (Δ = +3.2 W/kg, +18.4%, d = 0.76, p = 0.004). More importantly, FVimb decreased from 28.3% to 19.4% (absolute reduction –8.9 percentage points, relative reduction –31.4%, d = – 0.68, p = 0.024). CON showed no significant changes in any F-V variable.

**Table 3.**
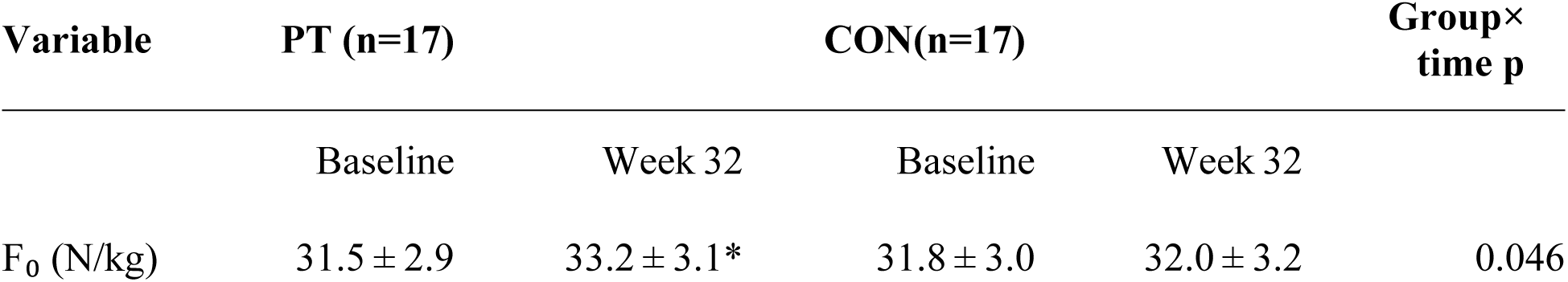
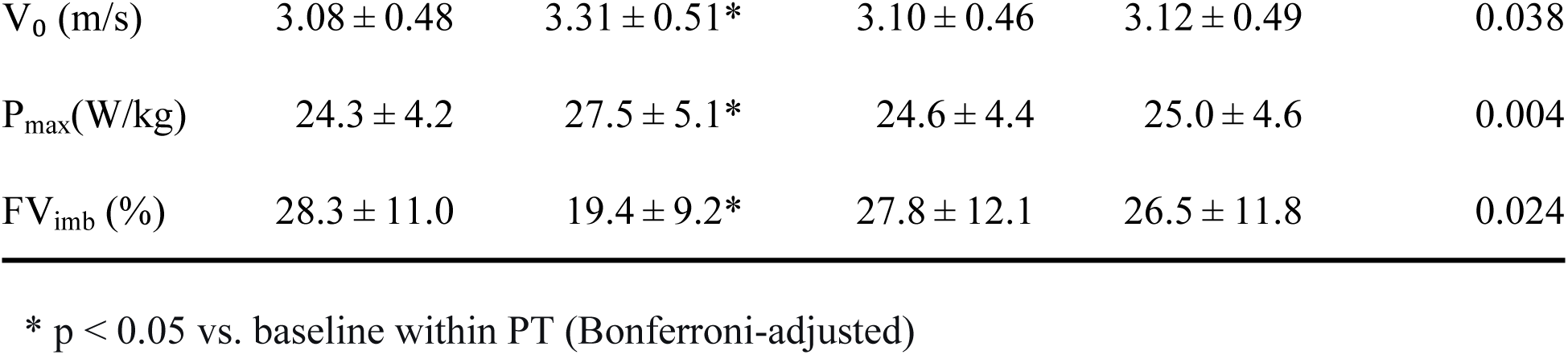
F-V variables at baseline and week 32.

### Correlations Between Changes

In the PT group (n=17), the reduction in FV_imb_ (ΔFV_imb_ = –8.9 ± 7.6%) correlated moderately with the gain in CMJ height (ΔCMJ = 5.3 ± 2.4 cm): r = –0.53, 95%CI [–0.78, – 0.15], p = 0.024 (**Figure 5A**). In contrast, ΔCMJ and ΔP_max_(3.2 ± 4.9 W/kg) showed a weak, non-significant relationship (r = 0.21, 95%CI [–0.28, 0.61], p = 0.39; **Figure 5B**). In Figure 5A, participants with baseline FVimb ≥28% (high imbalance) are marked as orange triangles; those with lower baseline imbalance appear as blue circles. The negative slope indicates that larger reductions in FVimb were associated with greater CMJ gains.

**Figure 5.**
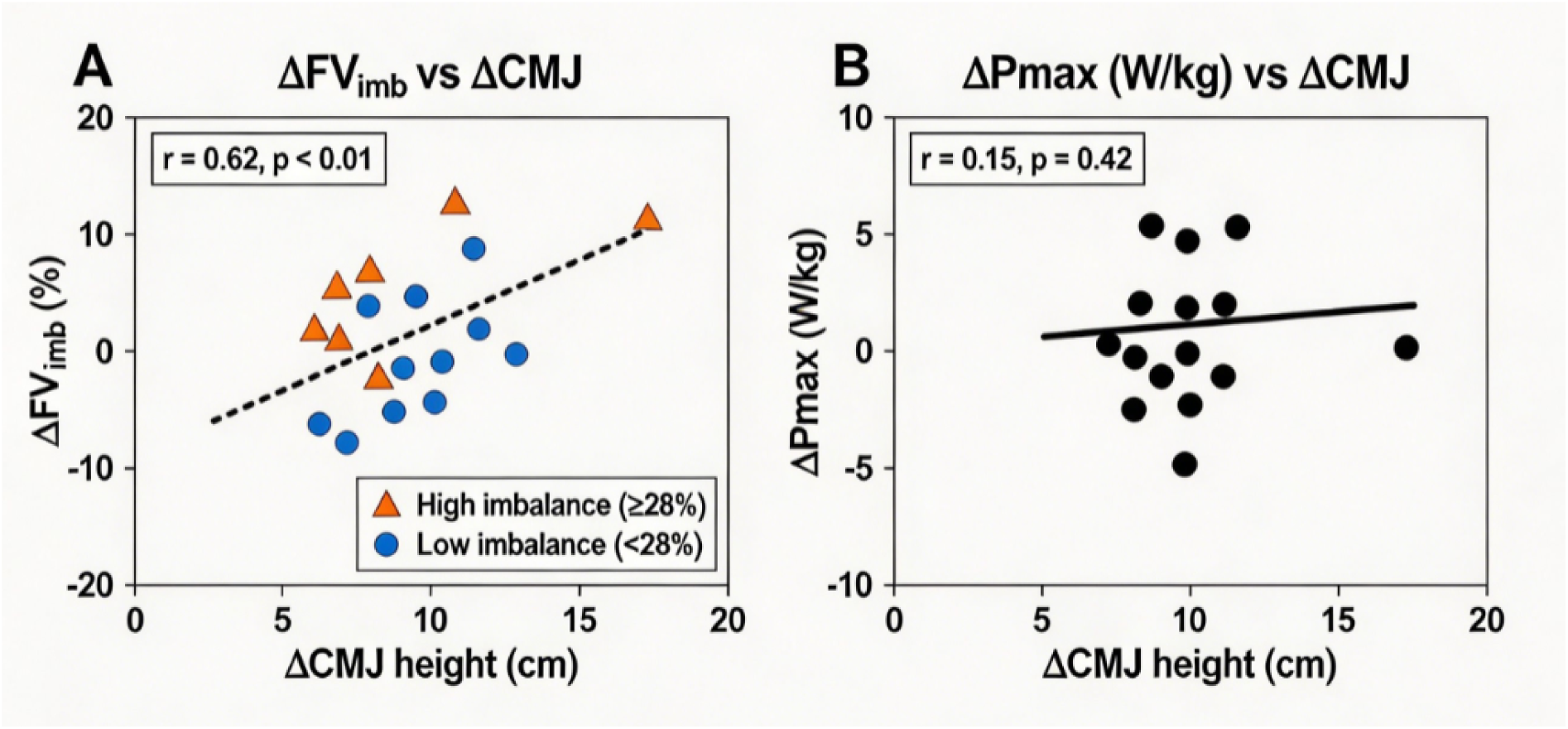
(A) Scatter plot of the change in CMJ height (ΔCMJ) versus the change in force-velocity imbalance (ΔFV_imb_) in the plyometric training group (n=17). Blue circles represent participants with baseline FV_imb_ <28%, orange triangles represent those with high imbalance (≥28%). The dashed line shows the linear fit (Pearson r = -0.53, 95%CI [-0.78, -0.15], p = 0.024). (B) Scatter plot of ΔCMJ versus change in maximal power (ΔP_max_) for the same participants. No significant correlation was observed (r = 0.21, 95%CI [-0.28, 0.61], p = 0.39). Blue circles: baseline FVimb < 28%; orange triangles: baseline FVimb ≥ 28%.

### Individual FV_imb_ Heatmap

**Figure 6** shows how individual FV_imb_(%) evolved over the five time points (baseline, weeks 8, 16, 24, 32) for the 17 PT participants. Rows are ordered from lowest baseline imbalance (bottom) to highest (top). The colour scale transitions from red (high imbalance, roughly ≥35%) through yellow to green (low imbalance, roughly ≤18%). The group average fell steadily from 28.3% at baseline to 19.4% by week 32. Most individuals followed a downward path, although with some spread – the cross-participant standard deviation shrank from 11.0% to 9.2%. Participants who started with the largest imbalances (top rows) often changed from red to yellow or green, meaning they experienced the biggest absolute improvements. Those already nearer the optimal range (bottom rows) changed only modestly. This visual evidence confirms that a non-individualised, periodized plyometric program can reduce F-V imbalance across the vast majority of athletes, with larger gains in those who have more room for improvement.

**Figure 6.**
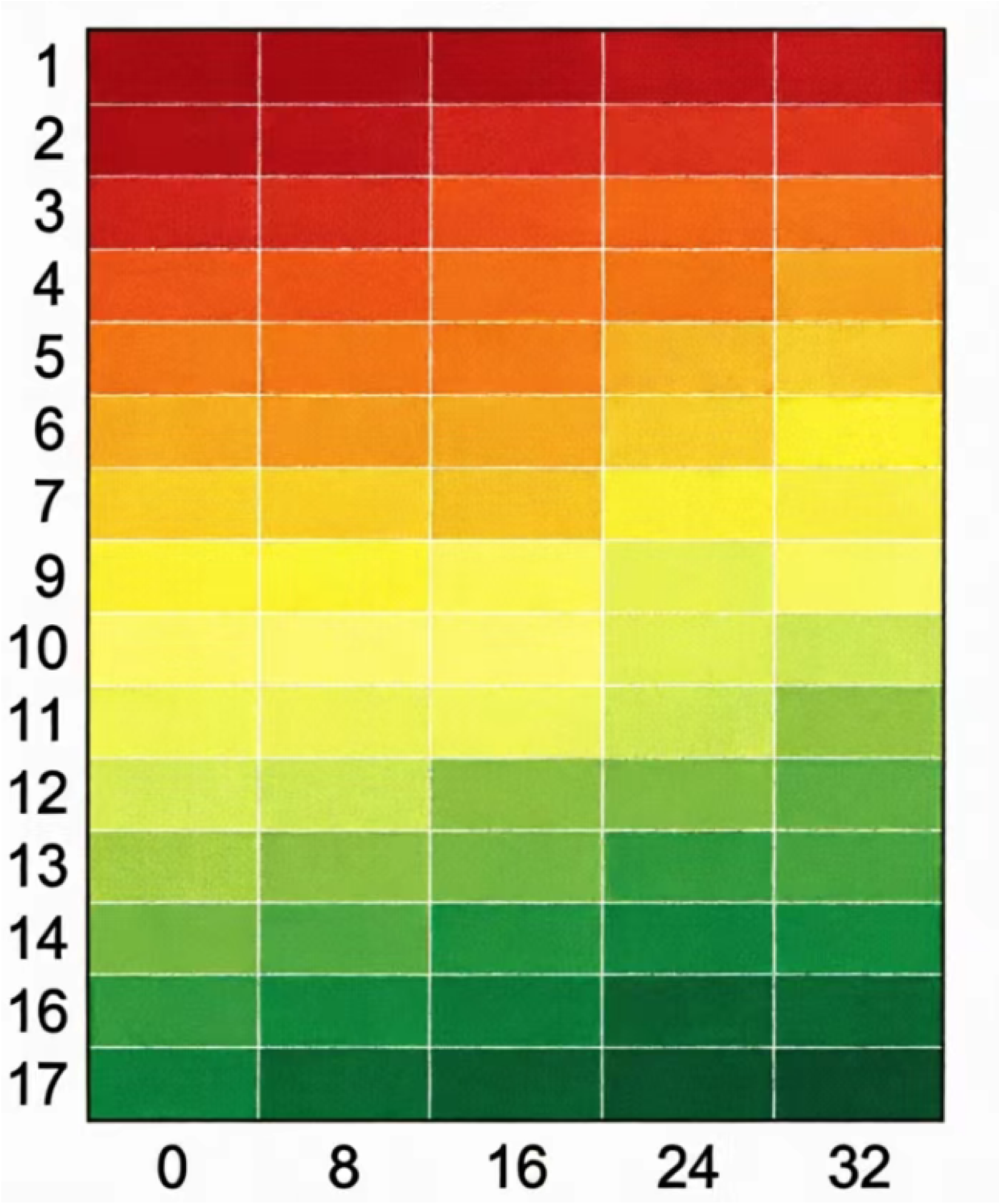
Heatmap of individual FV_imb_ (%) across 32 weeks (PT group, n=17). Rows sorted by baseline imbalance (bottom = lowest). Colour: red (high) → yellow → green (low). Group mean fell from 28.3% to 19.4%. Most rows shift from red/orange to green, showing that the periodized program improved F-V balance in nearly all participants, with the largest gains in those who started most imbalanced. Color scale: red (high imbalance) → yellow → green (low imbalance).

### Individual Force-Velocity Curve Shifts Toward Optimal

To illustrate the range of individual responses to the 32-week intervention, we selected three participants whose baseline FVimb values fell closest to the median of the high-imbalance (≥28%), moderate-imbalance (20–28%), and low-imbalance (<20%) tertiles, respectively. These cases represent a force-dominant, a balanced, and a velocity-dominant profile at baseline. Their F-V curves before and after training are shown in **Figure 7**, together with the theoretical optimal curve for a maximal power output of approximately 24.5 W/kg.

**Figure 7.**
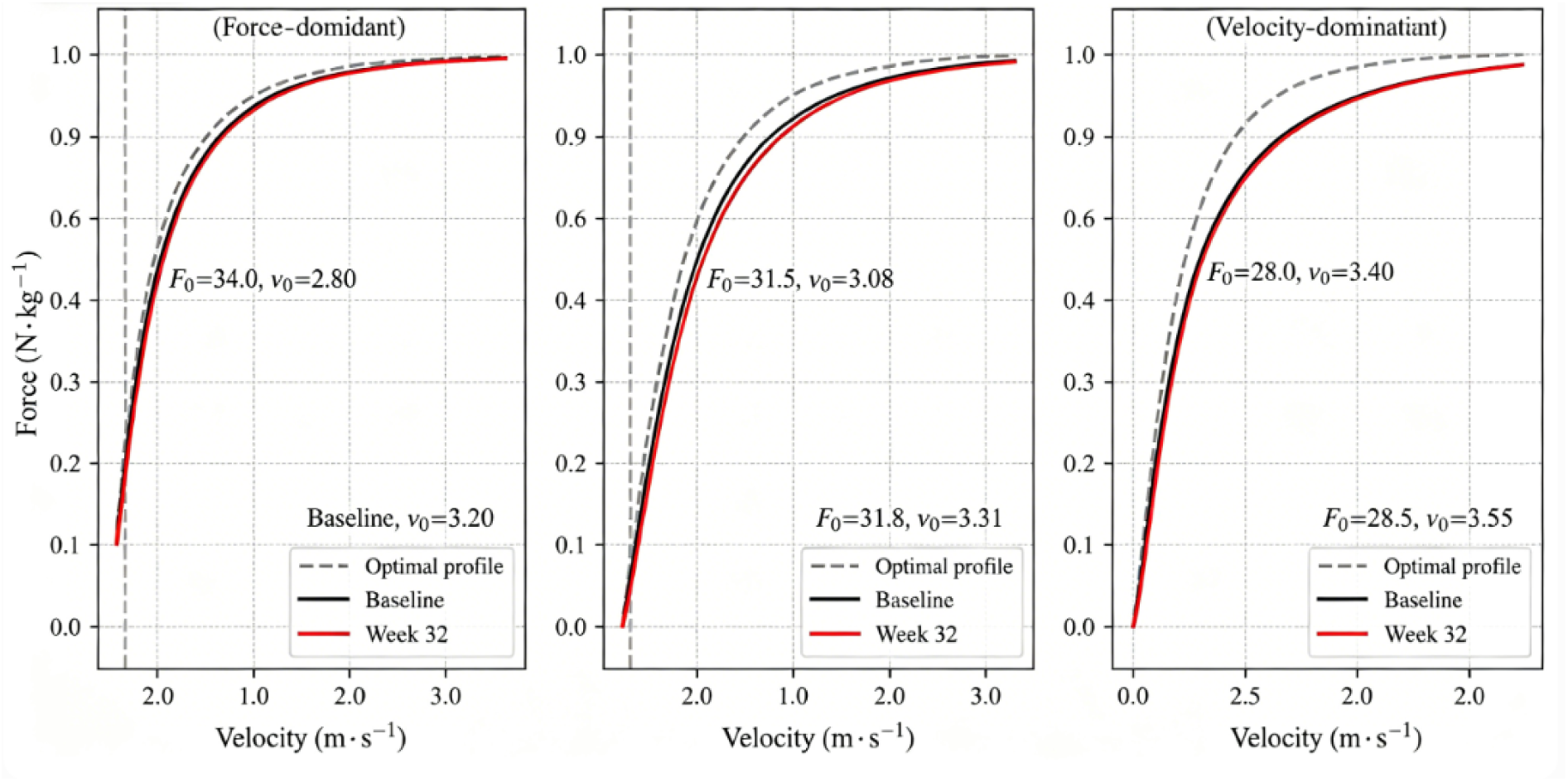
Individual force-velocity (F-V) curves at baseline and after 32 weeks for three representative participants from the PT group, compared with the theoretical optimal curve (dashed grey line). (A) High-imbalance (force-dominant) participant: baseline F₀ = 35.0 N·kg⁻¹, v₀ = 2.80 m·s⁻¹; post-training F₀ = 34.0 N·kg⁻¹, v₀ = 3.20 m·s⁻¹. (B) Moderate-imbalance participant: baseline 31.5 N·kg⁻¹, 3.08 m·s⁻¹; post 31.8 N·kg⁻¹, 3.31 m·s⁻¹. (C) Low-imbalance (velocity-dominant) participant: baseline 28.0 N·kg⁻¹, 3.40 m·s⁻¹; post 28.5 N·kg⁻¹, 3.50 m·s⁻¹. In each case, the curve moves closer to the optimal, indicating that the periodized program restores F-V coordination. Solid line = baseline; dashed line = post-training; grey dashed = theoretical optimal.

High-imbalance participant: At baseline, this player had a steep slope with F₀ = 35.0 N·kg⁻¹ and v₀ = 2.80 m·s⁻¹ – a pronounced force-dominant profile (slope –12.5 Ns·m⁻¹·kg⁻¹). After 32 weeks, F₀ decreased slightly to 34.0 N·kg⁻¹ (–2.9%), while v₀ increased markedly to 3.20 m·s⁻¹ (+14.3%). The slope flattened to –10.6 Ns·m⁻¹·kg⁻¹, moving considerably closer to the optimal curve. The absolute FV_imb_ reduction for this player was 18.5 percentage points. Moderate-imbalance participant: Baseline values (F₀ = 31.5 N·kg⁻¹, v₀ = 3.08 m·s⁻¹) were close to typical figures for collegiate volleyball players. The F-V curve already approximated the optimal shape. Following training, both F₀ (+1.0%) and v₀ (+7.5%) improved (to 31.8 N·kg⁻¹ and 3.31 m·s⁻¹). The curve shifted slightly outward and upward, resulting in an 8.9-percentage-point reduction in FV_imb_.

Low-imbalance participant: This player started with a shallow slope, characterised by relatively low F₀ (28.0 N·kg⁻¹) and high v₀ (3.40 m·s⁻¹) – a velocity-dominant pattern. By week 32, F₀ rose to 28.5 N·kg⁻¹ (+1.8%) and v₀ increased to 3.50 m·s⁻¹ (+2.9%). The slope became slightly steeper, moving closer to the optimal. FV_imb_ decreased by 3.4 percentage points.

Across these three cases, the direction of change consistently shifted the individual F-V curve toward the theoretical optimum: the force-dominant player sacrificed a small amount of force to gain substantial velocity; the velocity-dominant player added a little force while preserving high velocity; and the moderate player improved both dimensions. These qualitative observations align with the group-level finding that a periodized plyometric program reduces FV_imb_ across a wide range of initial profiles (Section 3.3).

## Discussion

This 32-week trial provides several novel observations regarding long-term plyometric training. A prolonged periodized program induced large, progressive improvements in CMJ height (5.3 cm, 16%) and reactive strength (31.6%). Notably, FV imbalance decreased markedly by 31.4% without any athlete-specific diagnosis. Furthermore, the gain in jump performance was associated with reduced FV imbalance rather than with maximal power alone, implying that restoring force-velocity coordination may play a central role.

### Long-Term Plyometric Effects: Beyond Short-Term Gains

The majority of published volleyball plyometric studies are limited to 12 weeks or fewer, documenting CMJ gains of just 5–10% [2,14]. Our 32-week program produced larger improvements than those shorter interventions. The 16% gain observed here after 32 weeks is considerably larger, but what stands out is the non-linear time course. The largest increments happened between weeks 8–16 and again between weeks 24–32, while weeks 16–24 showed a clear plateau (Table 2). This pattern aligns closely with the principles of periodization. The strength-power transition phase (weeks 9–16) might have improved the athletes’ ability to absorb and redirect force. That quality is often ignored in plyometric studies lasting ≤12 weeks. After that, the maximal power phase (weeks 17–24) likely induced considerable neuromuscular fatigue. Under those conditions, supercompensation typically requires a reduction in training load before it can manifest. Indeed, we only observed clear CMJ gains after we reduced volume in weeks 25–32. That supercompensation emerged only after the maintenance phase (weeks 25–32), suggesting that coaches who stop plyometrics after 16 weeks miss out on the delayed gains that come from unloading and recovery. In practical terms, the second half of a long-term program is not redundant; it is where the consolidation of earlier adaptations occurs.

Reactive strength index - modified (RSIₒd) improved by 31.6% in the plyometric group, an effect larger than the 10–20% changes reported in previous 6 - week studies [15]. The substantial improvement in RSImod (31.6%) suggests that the prolonged plyometric program enhanced the athletes’ ability to produce force rapidly during short ground contact tasks. While previous research has proposed that such improvements may be related to increased tendon stiffness and reduced neuromuscular delay within the stretch-shortening cycle [2], our study did not include direct measurements (e.g., muscle biopsies, electromyography, or ultrasound) to confirm these mechanisms. We therefore interpret the RSImod improvement as a performance-level adaptation without speculating on the underlying structural or neural contributors [16].

The earlier improvement in RSImod (by week 16) compared to CMJ height (which showed clear gains only after week 24) may indicate that the ability to produce force rapidly improves before this capacity translates into greater jump height. However, given the absence of direct mechanistic measurements, this temporal pattern should be interpreted descriptively. Alternative explanations include differences in the sensitivity of these two measures to training-induced changes, or differential rates of adaptation in the neural and muscular components of the stretch-shortening cycle. Our data do not allow us to distinguish between these possibilities.

For coaches, tracking RSI□ₒ_d_ on a weekly basis offers a practical way to spot positive neuromuscular adaptations weeks before they show up in standard jump tests.

### Remodelling the Force-Velocity Profile Without Individualisation

One ongoing debate in applied sport science concerns whether force-velocity profiling requires individualization to be practically useful. Jiménez-Reyes and colleagues [7] showed that 9 weeks of individually prescribed training based on F-V profiling reduced imbalance by approximately 40% and increased CMJ by 14%. In the present study, the 32-week generic program reduced FV imbalance by 31.4% and increased CMJ height by 16%. Although the magnitudes of improvement in both studies fall within a comparable range, direct numerical comparisons should be interpreted with caution due to differences in intervention duration, athlete level, testing protocols, and other methodological factors. The contribution of the present study lies in demonstrating that, in the absence of individual F-V diagnostics, a well-designed long-term periodized program can still produce meaningful remodelling of the F-V profile. Without a direct head-to-head trial comparing non-individualized versus individualized prescription, we cannot conclude equivalence or non-inferiority. Instead, our findings should be viewed as hypothesis-generating: a well-designed periodized program can produce substantial F-V adaptations, but whether it matches the efficiency of individualized diagnosis remains an open question.

At first glance, a one-size-fits-all training plan seems unlikely to drive consistent performance gains across athletes with varied physical profiles. Even so, the structured four-phase layout of the program clearly accounts for this positive outcome. Unlike programs that apply a uniform stimulus to all athletes, our periodized design rotated blocks emphasizing either heavy loading or high speed. That way, a force-deficient player got the heavy work they needed, while a velocity-deficient player got plenty of low-load explosive jumps – all without us having to test who was who. Across the 32 weeks, the program alternated between phases emphasizing force production (depth jumps, weighted squat jumps with 10% body mass) and phases emphasizing velocity and reactive ability (low-load reactive jumps, hurdle hops with minimal contact time). A player with a force-dominant profile (high F₀, low v₀) received ample high-speed work during the maintenance phase, while a player with a velocity-dominant profile (low F₀, high v₀) was exposed to heavy loading during the maximal power phase. A plausible—but untested—interpretation is that the periodized structure, by alternating between phases emphasizing force production and phases emphasizing velocity and reactive ability, may have provided varied stimuli that addressed different aspects of the F-V profile across the training cycle. However, our study did not measure individual responses to each training phase, nor did it compare outcomes against a non-periodized or individualized program. The mechanism underlying the observed F-V adaptations remains unknown. This does not negate the value of individualized training approaches. Instead, it indicates that for resource-constrained team sports, a standardized and properly designed general training plan can deliver the majority of performance benefits with far lower financial input and organizational workload.

Our exploratory subgroup analysis, while underpowered, added a nuance: players with baseline FV imbalance of 28% or higher (n=9) showed a non-significantly larger CMJ gain (6.2 cm) than those with lower imbalance (4.4 cm, p=0.11). Although the p-value did not reach conventional significance, the 1.8 cm difference in effect size is practically meaningful. This trend hints that athletes who are most imbalanced may have the most room for improvement and could benefit even more from the same program. Future studies should pre-specify subgroup analyses based on baseline imbalance tertiles or quartiles, with adequate statistical power to detect interaction effects. A 2025 meta-analysis by Wolte et al. [6] reached a conclusion similar to ours. According to their synthesis, individualized F-V profiling does not outperform well-designed non-individualized programs when it comes to performance gains.

### Why Does Reduced Imbalance Matter More Than Increased Pmax?

Many strength coaches view maximal power (Pmax) as the primary determinant of jump height. If peak power really drove jump height, we’d expect ΔCMJ and ΔPmax to track together. They didn’t. The correlation was weak (r=0.21) and not significant. Instead, ΔCMJ correlated moderately with the reduction in FV imbalance (r=-0.53, p=0.024). These findings align with the emerging “optimal F-V profile” concept [17], which argues that how force and velocity are balanced matters as much as—or more than—the absolute power output.

An athlete with a pronounced force-dominant profile – characterized by elevated F₀ alongside depressed v₀ – may still achieve a high Pmax. However, the majority of that power is generated at low movement speeds, making it poorly suited to an explosive, unloaded task such as a countermovement jump. The same applies to a velocity-dominant athlete: high v₀ but low F₀ means the power is delivered at velocities too high to generate sufficient impulse against body weight. In both cases, the mechanical effectiveness of the available power is compromised. Our training program simultaneously shifted F₀ upward by 5.4% and v₀ upward by 7.5%, effectively flattening the slope and bringing it closer to the theoretical optimum. This dual adjustment may explain why athletes jumped higher after training even when their Pmax did not increase dramatically. In essence, the same absolute power output becomes more useful because it is delivered at a more appropriate velocity for the task.

This insight has implications for how coaches interpret testing results. An athlete boasting greater Pmax alongside severe FV asymmetry often achieves poorer jumping capacity than those with marginally reduced peak power but well-aligned force–velocity balance. Training interventions that only aim to raise Pmax (e.g., maximal strength training alone) could actually worsen imbalance if they push F₀ up without improving v₀. Conversely, our periodized plyometric model, by alternating loading conditions, naturally improves both ends of the spectrum and restores coordination.

### Practical Implications for Collegiate Coaches

These findings offer several practical takeaways for college volleyball coaches. Long-term periodization matters. A 32-week block covering pre-season through post-season yields progressive, non-linear gains, whereas a single 8-week block—the typical duration found in the literature—is insufficient to fully reshape the force-velocity profile. Coaches should plan for at least 24 weeks of structured plyometrics to see substantial imbalance correction.

For programs that currently lack access to force-plate technology, our data demonstrate that a well-structured periodized model can still produce substantial FV imbalance reduction. However, this does not imply that force-plate testing is unnecessary or that individualized training is no better. Our study was not designed as an equivalence or non-inferiority trial against individualized F-V prescription. Therefore, we cannot recommend replacing individualized diagnostics with a generic program based on the present evidence. Instead, we suggest that when resources permit, individualized testing may offer additional benefits; conversely, when resources are limited, the present protocol represents a viable alternative that is superior to low-intensity control conditions. Direct comparisons between generic and individualized programs are needed before strong practice recommendations can be made. That said, if force plates are available, they can help identify athletes with extreme imbalance who might benefit from slight modifications (e.g., spending more time in the phase that targets their deficit).

A practical recommendation concerns routine monitoring of RSIₒd. It improved earlier (by week 16) than CMJ height (by week 24) and can be measured with nothing more than a contact mat or even a smartphone app. Weekly tracking of RSIₒd from a 30-cm drop jump provides an early window into neuromuscular adaptation, allowing coaches to adjust training loads before performance stagnates or declines.

### Limitations

Certain constraints in research methodology exist and need to be recognized when evaluating the present experimental results. A major constraint relates to our mechanistic inferences. We did not collect muscle biopsies or electromyographic data, which means our claims about neural versus structural adaptations—such as changes in motor unit recruitment, tendon stiffness, or muscle architecture—remain indirect. Direct evidence for these mechanisms would require additional measurements that were beyond the scope of this field-based study. Another limitation concerns the force-velocity profiling method itself. We used a two-point approach (unloaded and loaded squat jumps) following Samozino et al. [10], which assumes a strictly linear relationship between force and velocity. While this simplified method is practical for team settings, it may underestimate curvature in athletes with extreme profiles—for example, those who are exceptionally force-dominant or velocity-dominant. A multi-point method (e.g., three or four loads) would capture nonlinearities better but requires more testing time and equipment [18].

Long-term retention of the training effects remains unknown. We assessed outcomes immediately after the 32-week intervention but did not conduct a follow-up test at, say, 6 months after the program ended. Without such data, we cannot say whether the gains in jump height, reactive strength, and FV imbalance would persist, diminish, or even reverse once periodized plyometric training stops. Future studies should include delayed post-tests to answer this practical question.

The design of the control condition also introduces some ambiguity. Our active control group performed low-intensity conditioning (core stability, resistance bands, balance tasks) alongside their regular volleyball practice. However, they did not receive a sham high-load protocol that would perfectly mimic the time, effort, and perceived exertion of the plyometric program. Although total weekly training minutes were equated between groups (both performed 135–180 min of supplemental training per week), the lower mechanical demands of the CON condition mean we cannot completely rule out a dose-response effect. Nevertheless, the primary aim was to test the efficacy of a specific plyometric protocol against a typical low-intensity alternative—a comparison relevant to real-world coaching decisions.

Menstrual cycle phase was not standardized across testing sessions for female participants. Although the distribution of phases did not differ between groups, the lack of phase standardization may have increased within-subject variability and reduced the precision of our estimates in female athletes. Future studies should consider scheduling tests during a specific phase of the menstrual cycle (e.g., early follicular) to control for hormonal influences on neuromuscular performance.

Finally, our handling of missing data was adequate but not exhaustive. Only 2.8% of data points were missing, due to illness on scheduled test days. We used restricted maximum likelihood (REML) under the missing-at-random assumption, which is standard in mixed-model analyses. Nevertheless, we did not perform multiple imputation as a sensitivity check to confirm the robustness of our results. Doing so would have provided additional reassurance that the missing values did not bias the estimates.

None of these limitations undermine our core conclusions. However, we offer them as a roadmap for readers and future investigators: interpret our evidence with these caveats in mind, and design more definitive studies where possible.

### Conclusion

Thirty-two weeks of periodized plyometric training improved lower-limb power, reactive strength, and jumping performance, and reduced force-velocity imbalance by nearly one-third even without individual jump profiling. The correlation between improved jump performance and reduced imbalance – but not with increased maximal power alone – suggests that restoring coordination along the force-velocity spectrum is a key adaptive mechanism. These findings carry several practical implications for collegiate volleyball coaches. Plyometric training should not be confined to a short pre-season block but instead integrated into the annual training plan across a full 32-week cycle. Within the context of a collegiate team with limited access to individual F-V profiling, a well-designed periodized plyometric program delivered uniformly to the entire team effectively reshaped players’ F-V profiles and enhanced jumping performance relative to a low-intensity control. Whether this generic approach produces effects equivalent to individualized F-V training remains unknown, as the present study did not include an individualized prescription arm. Future head-to-head trials are required to determine if advanced laboratory testing provides additional benefits beyond a generic periodized program.

### Practical Applications

Based on the 32-week findings, collegiate volleyball coaches should consider the following practical strategies:

Adopt a season-long approach. Progressive gains in jump performance and force-velocity balance were not fully realized until after 24 weeks. A single 8-week pre-season block is insufficient; integrate plyometric training across the entire annual cycle.

Apply a four-phase periodized model. Follow a structured progression: anatomical adaptation (weeks 1–8, 80–120 low-impact contacts), strength-power transition (weeks 9–16, 100–150 contacts, 30–45 cm drops), maximal power (weeks 17–24, 60–80 contacts, 50–70 cm drops, 10% body mass load), and maintenance (weeks 25–32, 40–60 contacts, 2 sessions/week). This rotation naturally addresses both force-deficient and velocity-deficient profiles.

Monitor RSImod weekly. This reactive strength index improved earlier than CMJ height (week 16 vs. week 24) and requires only a contact mat or smartphone. Use it as an early indicator of positive neuromuscular adaptation to adjust training loads proactively.

Focus on force-velocity balance, not just peak power. Improvements in jump height were more closely related to reduced FV imbalance than to increased maximal power. When assessing training effectiveness, evaluate whether the athlete’s force-velocity profile is becoming better coordinated, not merely whether peak power has increased.

For teams lacking force-plate access, this standardized 32-week program provides a resource-efficient alternative that produced meaningful performance improvements in this cohort, requiring only basic equipment (hurdles, boxes, weighted vests). However, this does not rule out additional benefits from individualized F-V testing when resources permit.

## Ethics Statement

The study was approved by XYAFU Ethics Committee (XY-2024-029).

## Data Availability Statement

The datasets generated and analyzed during the current study are available from the corresponding author upon reasonable request.

## Acknowledgments

None.

## Author Contributions

H-MW: Conceptualization, Methodology, Investigation, Formal analysis, Writing – original draft. JL: Conceptualization, Resources, Supervision, Funding acquisition, Writing – review & editing. All authors contributed to the article and approved the submitted version.

---

**Supplementary Table S1.** Monitoring of training load, match exposure, and potential confounders over 32 weeks (mean ± SD or median [IQR] as appropriate)

| Variable | PT group (n=17) | CON group (n=17) | Between-group p-value | Notes |
| --- | --- | --- | --- | --- |
| **External training load** |  |  |  |  |
| Volleyball technical/tactical training (h·week<sup>-1</sup>) | 8.4 ± 1.2 | 8.3 ± 1.3 | 0.82 | Self-reported team log |
| Matches played (sets·week<sup>-1</sup>) | 4.2 [3.0, 5.5] | 4.0 [2.5, 5.0] | 0.67 | Regular season (weeks 9–24) |
| **Internal training load** |  |  |  |  |
| Session-RPE (0–10, PT sessions only) | 6.3 ± 1.1 | — | — | For CON: low-intensity conditioning (RPE 2–3) |
| Session-RPE (0–10, CON sessions only) | — | 2.5 ± 0.7 | — | Between-group comparison not applicable due to different content |
| **Additional resistance training (h·week<sup>-1</sup>)** | 0.0 (0.0–0.0) | 0.0 (0.0–0.0) | 0.99 | Prohibited per protocol; compliance verified weekly |
| **Injury/illness episodes** |  |  |  |  |
| Total injuries (n) | 2 | 3 | — | One ankle sprain (PT) during match; none related to intervention |
| Total illness days missed | 4 | 6 | — | Common cold, gastrointestinal |
| **Female participants only (n=9 per group)** |  |  |  |  |
| Menstrual cycle phase at testing (baseline, week 8, 16, 24, 32) |  |  |  |  |
| - Early follicular (days 1–5) | 18 / 45 tests | 20 / 45 tests | 0.78 | Self-reported |
| - Late follicular (days 6–14) | 12 / 45 | 10 / 45 | 0.82 |  |
| - Luteal (days 15–28) | 15 / 45 | 15 / 45 | 1.00 |  |
| Oral contraceptive use (n users) | 4 | 5 | 0.71 | Stable throughout trial |
| **Lifestyle factors** |  |  |  |  |
| Sleep duration (h·night<sup>-1</sup>) | 7.2 ± 0.8 | 7.1 ± 0.9 | 0.79 | Weekly diary |
| Subjective sleep quality (1–5, higher = better) | 3.8 ± 0.6 | 3.7 ± 0.7 | 0.65 | |

| Variable | PT<br>group<br>(n=17) | CON<br>group<br>(n=17) | Between-group<br>p-value | Notes |
| --- | --- | --- | --- | --- |
| Nutritional supplement use<br>(n users) | 3 (whey<br>protein) | 2 (vitamin<br>D) | 0.63 | No ergogenic supplements<br>allowed |
| **Season timing** |  |  |  |  |
| Pre-season (weeks 1–8) | — | — | — | Lower match volume |
| In-season (weeks 9–24) | — | — | — | High match volume (2–3<br>matches/week) |
| Post-season (weeks 25–32) | — | — | — | Reduced match volume |

**Abbreviations:** PT, plyometric training group; CON, control group; RPE, rating of perceived exertion; h, hours; min, minutes.
**Notes:** Data are presented as mean ± SD for normally distributed variables, or median [interquartile range] for non-normally distributed variables.
Menstrual cycle phase was self-reported; tests were not standardized for phase, but distribution across phases did not differ between groups ( $p > 0.05$ for all comparisons).
Sleep and nutritional data are based on weekly questionnaires; no objective monitoring (e.g., actigraphy) was performed.

**Supplementary Table S2.** Mixed-model ANOVA results for CMJ height with sex as a covariate.

| Effect | F-value | df | p-value | Partial $\eta^2$ |
| --- | --- | --- | --- | --- |
| Group | 6.82 | 1, 31 | 0.014 | 0.180 |
| Time | 28.45 | 4, 124 | <0.001 | 0.479 |
| Sex | 42.17 | 1, 31 | <0.001 | 0.576 |
| Group × Time | 9.73 | 4, 124 | <0.001 | 0.239 |
| Group × Sex | 0.52 | 1, 31 | 0.477 | 0.016 |
| Time × Sex | 1.38 | 4, 124 | 0.245 | 0.043 |
| Group × Time × Sex | 1.12 | 4, 124 | 0.349 | 0.035 |

**Note:** After adjusting for sex, the Group × Time interaction remained significant ( $p < 0.001$ ), confirming that the plyometric intervention improved CMJ height independently of sex. No three-way interaction was observed, indicating that the training effect did not significantly differ between male and female athletes.

**Supplementary Table S3.** Estimated weekly jump volume during regular volleyball technical/tactical drills (mean ± SD).

| Phase | PT Group<br>(jumps/week) | CON Group<br>(jumps/week) | Between-group<br>p-value |
| --- | --- | --- | --- |
| Baseline (Pre-intervention) | 142 ± 31 | 138 ± 28 | 0.69 |
| Phase I (Weeks 1–8) | 135 ± 29 | 140 ± 33 | 0.64 |
| Phase II (Weeks 9–16) | 148 ± 35 | 145 ± 30 | 0.78 |
| Phase III (Weeks 17–24) | 155 ± 38 | 150 ± 35 | 0.68 |
| Phase IV (Weeks 25–32) | 130 ± 26 | 128 ± 24 | 0.82 |

Note: Jump counts were estimated from video analysis of two randomly selected technical sessions per week (spiking and blocking drills). Only jumps >20 cm in estimated height were counted. No significant between-group differences were observed at any phase (all $p > 0.05$ ), confirming that the additional plyometric load from the intervention was the primary differential stimulus between groups.

**Supplementary Figure S1.**
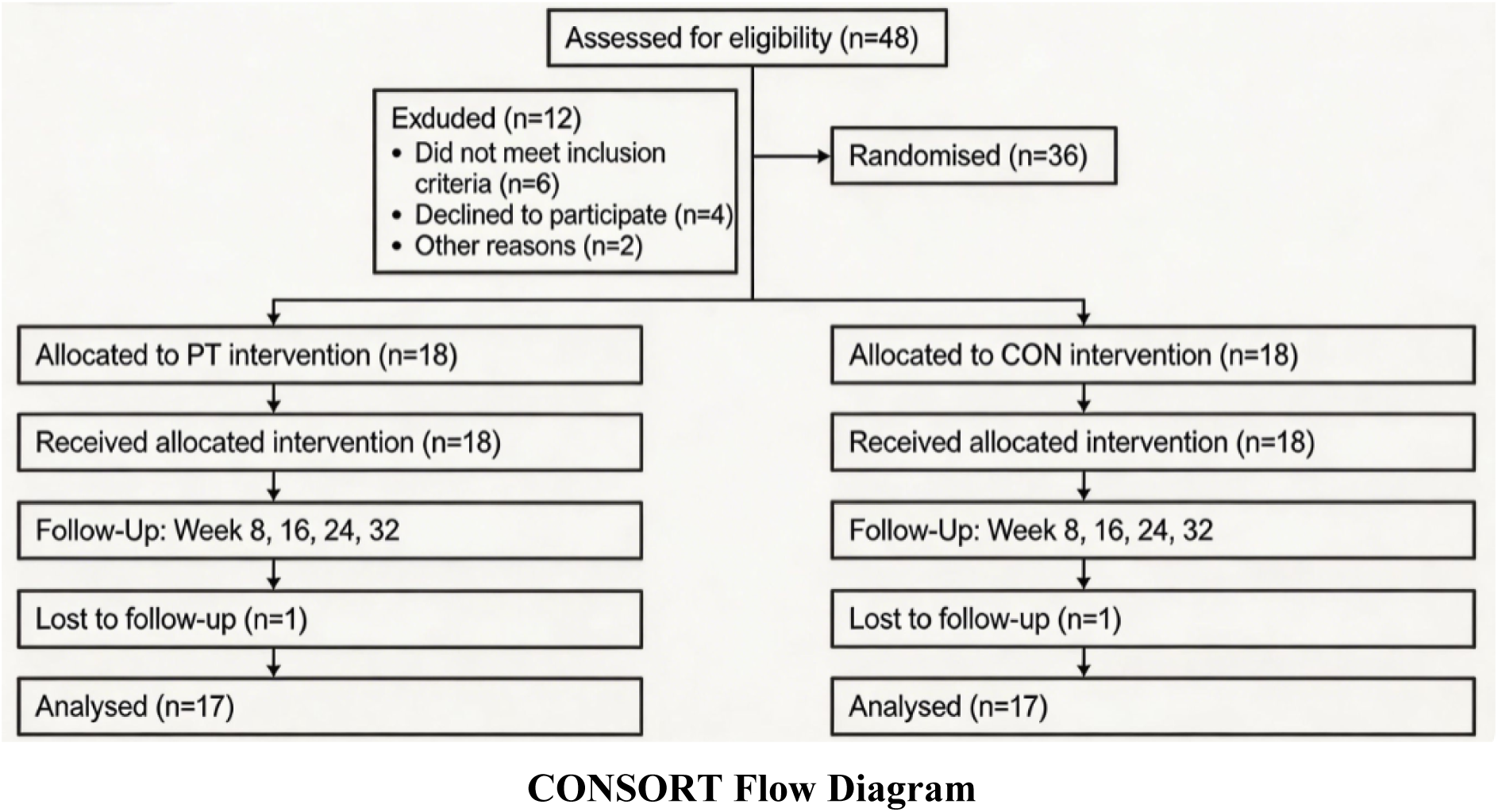
CONSORT 2010 flow diagram. Participant recruitment, randomisation, allocation, follow-up, and analysis for the 32-week randomized controlled trial. PT = plyometric training group; CON = control group.

**Supplementary Figure S2.**
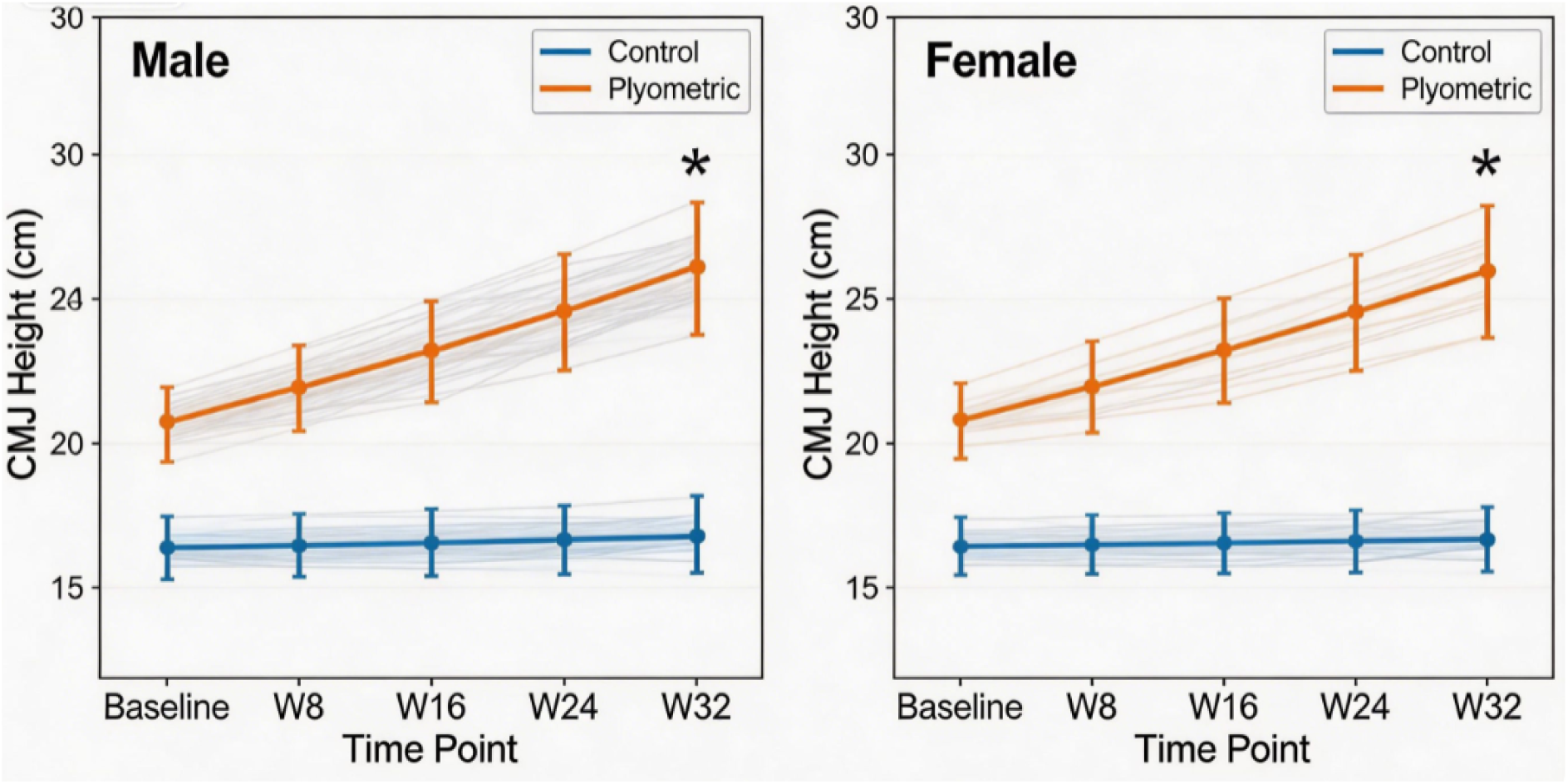
Changes in countermovement jump (CMJ) height over 32 weeks, stratified by sex. (A) Male athletes (PT: n=9; CON: n=8). (B) Female athletes (PT: n=8; CON: n=9). Data are presented as mean ± SD. Both sexes showed progressive improvements in the PT group, while CON groups remained relatively stable. The absence of a significant Group × Time × Sex interaction (p = 0.349) indicates that the training effect was consistent across sexes.

## Supplementary Methods: Force-Velocity Profiling Calculations

## 1. Two-point method (Samozino et al., 2008)

The force-velocity (F-V) relationship was modelled using the two-point method based on squat jumps (SJ) performed under two conditions: unloaded (body mass, BM) and loaded (BM + 20% BM via weighted vest).

For each condition, jump height (h) was derived from flight time (*t_f_*) using:

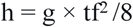

where g=9.81 m⋅s^−2^.

The push-off distance (*d_push_*) was estimated from the athlete’s body height (*H*) using the individualized regression equation (Samozino et al., 2008):

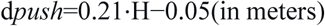

For each load condition, the theoretical maximal velocity (*v*) at take-off was:

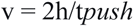

where t_push_ is the push-off duration (from the start of upward movement to take-off), and mean propulsive force (*F*) was:

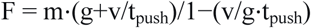

with *m* = system mass (body mass + load).

## 2. Linear regression and F-V variables

The F-V relationship was fitted with a linear regression:

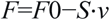

where:

F0= theoretical maximal force (intercept, N·kg⁻¹),

*v*0= theoretical maximal velocity (x-intercept, m·s⁻¹),

*S* = slope of the F-V relationship.

Maximal power (P_max_, W·kg⁻¹) was computed as:

Pmax=F0⋅v0/4

## 3. Force-velocity imbalance (FV_imb_)

Following Morin and Samozino (2016), the optimal velocity (vopt) for a given P_max_ was calculated as:

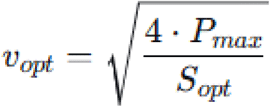

where *Sopt* is the optimal slope that satisfies:

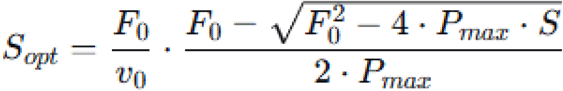

(Simplified implementation in practice: vopt is derived iteratively from the condition where the F-V slope equals the theoretical optimal slope for the measured Pmax.)

The imbalance index was then:

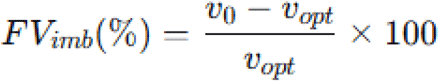

A positive FV_imb_indicates a velocity deficit (slope too steep, force-dominant); a negative value indicates a force deficit (slope too shallow, velocity-dominant).

## 4. Theoretical optimal curve for **Figure 7**

For each representative participant, the theoretical optimal F-V curve (dashed grey line in Figure 7) was constructed as follows:

Using the participant’s post-training Pmax (Week 32), the optimal force (Fopt) and optimal velocity (vopt) were determined from:

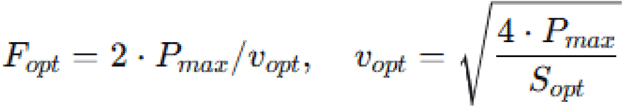

The curve was plotted as a straight line from *Fopt*on the y-axis to *vopt*on the x-axis, representing the ideal F-V relationship for that specific power output (i.e., the curve that maximises jump height for that *Pmax*).

The line equation for the optimal curve is:

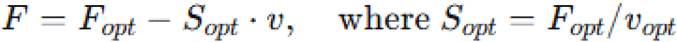

This visual comparison allows the reader to assess how each participant’s actual F-V slope (pre- and post-training) converged towards the theoretical optimum.

